# AI-Guided Multi-Objective Engineering of Glucoamylase Enables Acidification-Free Starch Saccharification

**DOI:** 10.64898/2026.09.24.753989

**Authors:** Jie Qiao, Xiaoru Ma, Yibo Song, Qiufeng Deng, Xinyue Ni, Guodong Liu, Shuaiqi Meng, Feihong Shi, Likang Deng, Haiyang Cui, Xiujuan Li

## Abstract

Glucoamylase is essential for industrial starch saccharification, but the limited thermostability and near-neutral pH tolerance of fungal glucoamylases necessitate cooling and acidification of liquefied starch. Here, we developed an artificial intelligence-guided strategy to simultaneously improve the thermostability, pH tolerance, and catalytic activity of glucoamylase from Penicillium oxalicum (PoGA). Two property-specific machine-learning models, CASPE-T and CASPE-A, identified substitutions associated with thermostability and pH tolerance, respectively. Experimental screening identified beneficial substitutions in 11 of 21 CASPE-T and 12 of 22 CASPE-A candidates. Folding-energy-guided recombination integrated the two traits while maintaining structural compatibility. The optimal variant, PoGA T513E/Q305N, exhibited 2.21-fold higher specific activity than the wild type, with half-life extended from 22.3 to 57.9 min at 60 °C and from 16.6 to 64.7 min at pH 8.0. Molecular dynamics simulations attributed these improvements to reinforcement of high-occupancy hydrogen-bonding networks, suppression of conformational fluctuations in the linker and carbohydrate-binding module, enhanced long-range dynamic coordination, and preservation of a compact catalytic architecture. At 60 °C and pH 6.5 without acidification, PoGA T513E/Q305N produced 219.9 g/L glucose and achieved 89.1% starch conversion, 31.4% higher than the wild type. This work provides an efficient framework for multi-objective enzyme engineering and sustainable starch biorefining.

**Entry for the Table of Contents:** We present a computational workflow combining property specific machine learning with folding energy guided recombination to achieve simultaneous multi property optimization. By efficiently navigating vast sequence spaces, this rational design pipeline successfully tailors biocatalysts for stringent manufacturing realities, ultimately enabling a completely streamlined industrial saccharification process that bypasses bulk acid addition.

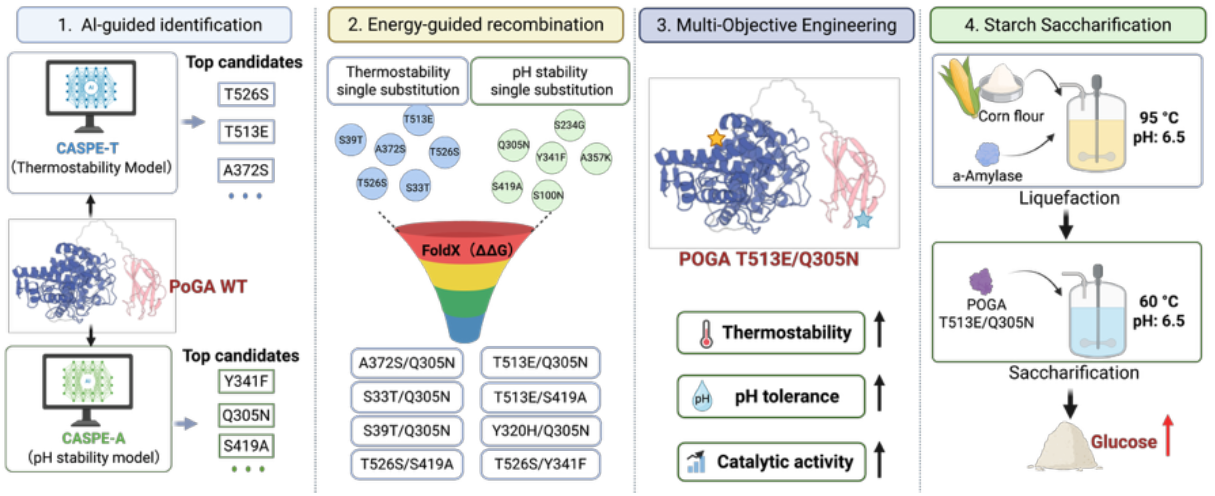

## Introduction

Starch is one of the most abundant renewable carbon resources and serves as the primary feedstock for the production of glucose syrup, high-fructose syrup, fuel ethanol, organic acids, and numerous bio-based chemicals ^1–5^. Efficient enzymatic saccharification of liquefied starch by glucoamylase (GA, EC 3.2.1.3) is therefore a cornerstone of modern starch biorefineries ^6, 7^. Compared with acid hydrolysis, enzymatic saccharification offers superior substrate specificity, higher glucose yields, lower energy consumption, and reduced by-product formation ^8^. Consequently, glucoamylases have become among the most extensively used industrial enzymes ^9^.

Despite their widespread application, the operational robustness of naturally occurring glucoamylases remains a major limitation for industrial starch processing ^6, 10^. Commercial starch conversion typically consists of high-temperature liquefaction of the raw material using α-amylase at approximately 85–105 °C and pH 6.0–6.5, followed by glucoamylase-catalyzed saccharification ^11^. Because most fungal glucoamylases exhibit optimal activity under acidic conditions (pH 4.0–5.0) and rapidly lose activity at near-neutral pH, the liquefied starch slurry must first be cooled and acidified before saccharification ^6, 10, 12^. These additional processing steps increase chemical consumption, energy demand, equipment corrosion, and production costs, ultimately reducing process sustainability. Therefore, developing glucoamylases capable of maintaining high catalytic efficiency under elevated temperatures and near-neutral pH is highly desirable for next-generation starch biorefineries.

Protein engineering has emerged as a powerful strategy for improving enzyme performance ^13, 14^. Directed evolution, rational design, and computational approaches have successfully enhanced the thermostability, catalytic activity, or substrate specificity of numerous industrial enzymes ^15–19^. For example, integrating bioinformatics mining with computational protein design identified and optimized a novel glucoamylase, resulting in a mutant with a 21% increase in specific activity and a 25% longer half-life at 40 °C ^20^. Furthermore, domain shuffling between two fungal glucoamylases generated a chimeric enzyme with enhanced thermostability, increasing the residual activity after incubation at 50 °C from 73% to 90% ^21^. More recently, rational engineering by introducing disulfide bonds and optimizing charge–charge interactions produced a glucoamylase variant with a 10 °C increase in optimum temperature and a 2.2-fold increase in specific activity ^11^. However, simultaneous optimization of multiple industrially relevant properties remains challenging because thermostability, pH tolerance, and catalytic efficiency are governed by highly coupled structural networks. Substitution that improves one property frequently compromises another, while the vast combinatorial sequence space makes experimental exploration prohibitively expensive. Although recent advances in machine learning have significantly accelerated enzyme engineering, most current models are optimized for a single objective and rarely address the simultaneous improvement of multiple environmental adaptation traits ^22^.

Here, we present an artificial intelligence-guided protein engineering strategy for the simultaneous enhancement of the thermostability and alkaline tolerance of the glucoamylase from *Penicillium oxalicum* (PoGA). Two specialized machine-learning models called CASPE-T and CASPE-A were employed to independently identify substitutions associated with thermal stability and pH tolerance ^23^. CASPE (Critical Amino Acids Streamline Protein Evolution) was developed as a lightweight protein-engineering platform for the precise identification and optimization of functionally critical residues. CASPE comprises two complementary modules: CAS (Critical Amino Acid Sites) and APCNet (Amino Acid Point Cloud Classification Network). CAS integrates gradient-weighted class activation mapping with multilayer attention matrices to extract property-relevant information directly from protein language models and translate it into interpretable residue-level importance scores, without requiring structural information or prior knowledge. Subsequently, the identified beneficial substitutions from CASPE were subsequently integrated through ΔΔG-guided recombination to maximize synergistic effects while preserving structural stability. The resulting variants exhibited substantially improved thermostability, alkaline resistance, and catalytic activity. Molecular dynamics simulations further revealed that enhanced robustness originated from reinforced hydrogen-bonding networks, improved long-range dynamic coupling, and stabilization of the catalytic architecture. Finally, industrial saccharification experiments demonstrated that the engineered enzymes maintained high glucose productivity under liquefaction-compatible conditions (60 °C and pH 6.5), enabling saccharification without prior pH adjustment and further cooling. This work establishes an efficient AI-assisted framework for multi-objective enzyme engineering and provides robust glucoamylase biocatalysts for more sustainable starch biorefining.

## Results and Discussion

### Biochemical and structural characterization of PoGA reveals limited thermostability and pH tolerance

As shown in **Fig. 1a and b**, biochemical characterization of PoGA using soluble starch as a substrate revealed exhibited optimal activity at pH 4.5 and 60°C, consistent with most fungal glucoamylases ^20, 34^. However, PoGA displayed a marked loss of stability under near-neutral and alkaline conditions. Specifically, the residual activity dropped sharply from 70.13% at pH 6.0 to 42.38% at pH 6.5 after incubation (**Fig. 1c**), indicating that even a modest increase in pH severely compromises enzyme stability. Similarly, thermostability assays demonstrated its severe vulnerability to heat stress, with residual activity plunging to 32.95% after incubation at the temperature of 60 °C (**Fig. 1d**). This dual sensitivity to both pH and thermal stress limits the direct application of PoGA in industrial starch processing, where liquefaction streams typically exit at an elevated pH of approximately 6.5 and high temperatures ^35^. Therefore, PoGA represents an ideal model enzyme for addressing the long-standing challenge of simultaneously improving thermostability and pH tolerance in industrial glucoamylases.

**Fig. 1.**
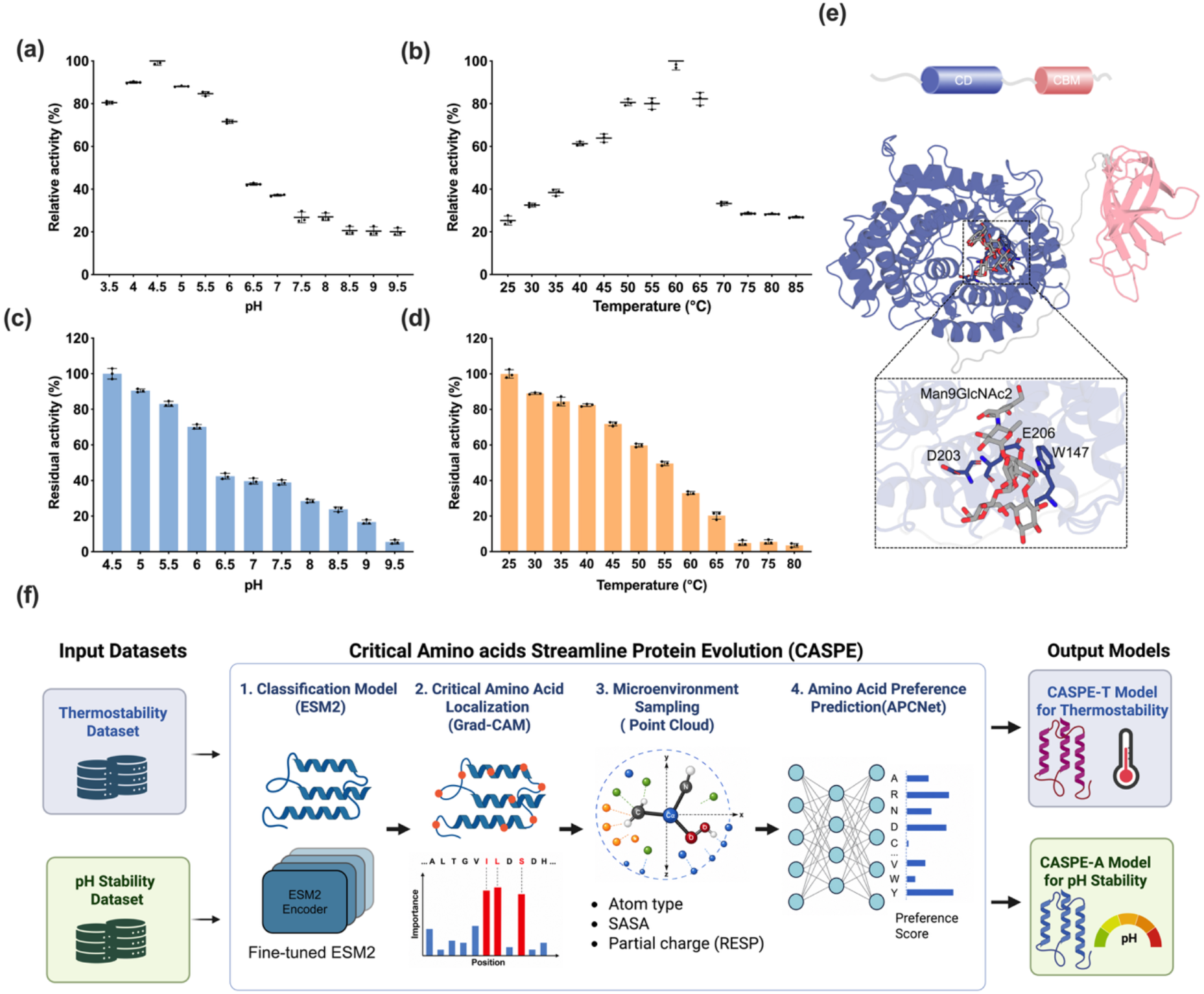
Biochemical properties and structure of PoGA WT, and an overview of the CASPE machine learning framework. (a) The pH-activity profile of PoGA. The enzymatic activity was measured at various pH levels ranging from 3.5 to 9.5; (b) The temperature-activity profile of PoGA. The optimal temperature was determined by assaying activity between 25°C and 85°C; (d) pH stability of PoGA. Residual activity was measured after incubating the enzyme in buffers of different pH values for 30 min. (e) Thermostability profile of PoGA. Residual activity was determined after incubating the enzyme at indicated temperatures for 30 min. (e) Structural model of PoGA: catalytic domain (CD, blue), carbohydrate-binding module (CBM, pink), and grey linker, D203 and E206 (active sites) and W147 (binding site); (f) Overview of the Critical Amino acids Streamline Protein Evolution (CASPE) framework. Protein sequences from thermostability and pH stability datasets were used to fine-tune an ESM2 classifier. Critical residues were identified by Grad-CAM, and local microenvironment features were analyzed by APCNet to predict optimal amino acid substitutions. The resulting CASPE-T and CASPE-A models guided thermostability- and pH stability-directed enzyme engineering

To establish a structural framework for subsequent engineering, we analyzed the three-dimensional architecture of PoGA. The predicted structure exhibits the canonical glucoamylase-fold, consisting of an N-terminal (α/α)6-barrel catalytic domain connected to a C-terminal carbohydrate-binding module (CBM) through a long, flexible linker (**Fig. 1e**) ^36, 37^. Molecular docking using Man9GlcNAc2 as a model substrate revealed a spacious substrate-binding pocket capable of accommodating bulky branched glycans. The catalytic residues D203 and E206 form the conserved catalytic acid/base pair responsible for glycosidic bond cleavage ^38^, whereas W147 serves as a key substrate-recognition residue by stabilizing the substrate through aromatic stacking and reinforcing the catalytic configuration of E206 via hydrogen-bond interactions ^39, 40^. These structural features establish the molecular basis for catalysis and provide a rational foundation for subsequent machine learning-guided protein engineering.

To facilitate rational enzyme engineering, we adopted the CASPE framework (**Fig. 1f**). CASPE is a machine learning-based protein engineering platform that predicts the fitness effects of amino acid substitutions by integrating sequence-derived evolutionary information with local structural microenvironment features under a PointNet architecture, thereby enabling efficient prioritization of beneficial substitutions while substantially reducing the experimental screening burden. Protein sequences from thermostability and pH stability datasets were used to construct the CASPE-T and CASPE-A models, respectively. Briefly, critical residues were identified using a fine-tuned ESM2 classifier coupled with Gradient-weighted Class Activation Mapping (Grad-CAM), and the optimal amino acid substitutions were subsequently predicted by the **Amino** Acid Preference Classification Network (APCNet) based on local three-dimensional microenvironment features. ^23^

### AI-guided identification of thermostability- and pH tolerance-determining substitutions

To identify substitutions associated with enhanced thermostability and pH stability, the CASPE-T and CASPE-A models were applied to predict the mutational fitness landscape of PoGA. The resulting fitness maps revealed distinct substitution preferences for the two target properties (**Fig. 2a, b**). Based on the predicted fitness scores, the top 21 single-site substitutions across distinct amino acid positions were selected from the CASPE-T model for experimental evaluation (**Fig. 2c**), whereas the CASPE-A model identified 22 unique candidate substitutions were predicted to enhance pH tolerance (**Fig. 2d**). Interestingly, the two fitness landscapes exhibited markedly different hotspot distributions, with only limited overlap between the residues prioritized by the two models. These results indicate that the molecular determinants governing thermostability and alkaline tolerance are largely distinct and cannot be captured by a single generalized prediction model.

**Fig. 2.**
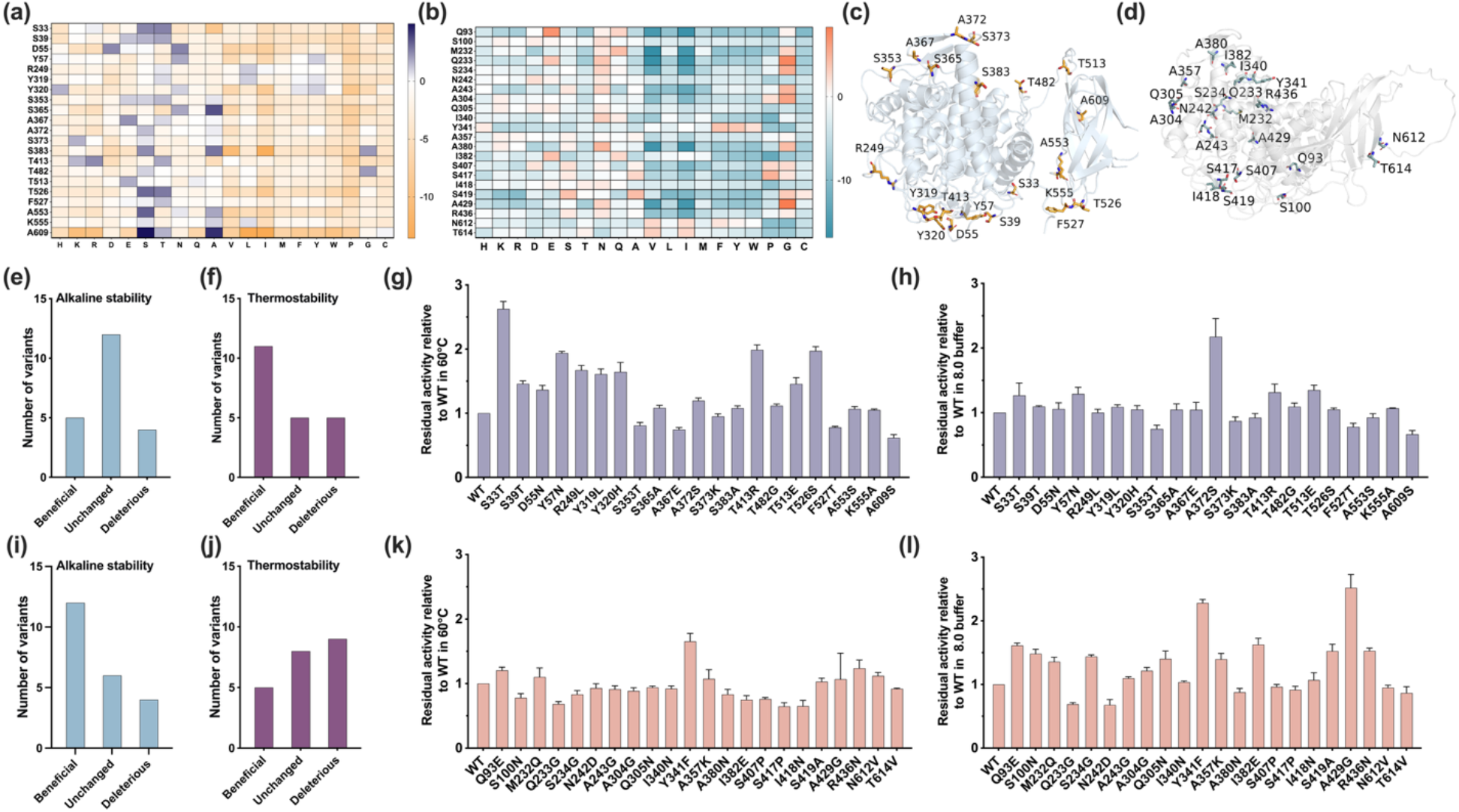
Computational screening and experimental validation of PoGA variants using the CASPE framework. Heatmaps showing the predicted fitness landscapes of single site mutations generated by the thermostability oriented CASPE-T model (a) and the alkaline stability oriented CASPE-A model (b). Structural mapping of the predicted candidate residues on the PoGA structure for the CASPE-T (c) and CASPE-A (d) models. (e-h) Experimental characterization and validation summary of the selected CASPE-T variants. Summary distributions showing the number of variants classified as beneficial, unchanged, or deleterious for alkaline stability (e) and thermostability (f). Residual activities relative to the WT were measured after incubation at 60 °C (g) and in pH 8.0 buffer (h). (i-l) Experimental characterization and validation summary of the selected CASPE-A variants. Summary distributions showing the number of variants classified as beneficial, unchanged, or deleterious for alkaline stability (i) and thermostability (j). Residual activities relative to the WT were measured after incubation at 60 °C (k) and in pH 8.0 buffer (l).

Experimental validation confirmed the high predictive accuracy of the CASPE-T model for thermostability engineering. Among the 21 predicted variants, 11 (52.38%) exhibited enhanced thermostability (>1.1-fold residual activity relative to the WT after thermal treatment), whereas five variants showed no significant change (0.9–1.1-fold) (**Fig. 2f**). This hit rate greatly exceeds that typically achieved by conventional directed evolution (~1%) ^41^, highlighting the efficiency of AI-guided mutation prioritization. Among the beneficial variants, S33T exhibited the greatest improvement, retaining 2.63-fold higher residual activity after incubation at 60 °C, while T482G and T526S also improved thermostability by nearly twofold (**Fig. 2g**). A similarly high prediction accuracy was obtained for the CASPE-A model. Of the 22 prioritized variants, 12 (54.55%) displayed enhanced alkaline stability (**Fig. 2i**), with A429G and Y341F exhibiting 2.51-fold and 2.30-fold higher residual activity at pH 8.0, respectively (**Fig. 2l**).

Interestingly, several thermostability-enhancing mutations also conferred improved alkaline tolerance. In particular, 23.81% of the CASPE-T variants exhibited enhanced stability at pH 8.0, with A372S showing the largest improvement (2.18-fold; **Fig. 2e, h**). Besides, five variants (22.73%) obtained from CASPE-A also showed improved thermostability (**Fig. 2j**), among which Y341F retained 1.65-fold higher residual activity after incubation at 60 °C (**Fig. 2k**). Although the two models identified largely distinct mutational hotspots, the cross-improvement observed in a subset of variants suggests that thermostability and alkaline tolerance could be mostly governed by shared structural determinants, providing a molecular basis for subsequent recombination engineering.

### Folding energy-guided recombination maximize the synergistic effect between thermostability and alkaline tolerance

To obtain PoGA variants with simultaneously improved thermostability and pH tolerance, we implemented a combinatorial recombination strategy by leveraging the beneficial single-point subsitution identified from the dual CASPE libraries. This approach aimed to explore the potential synergistic effects arising from the coupling of superior substitution sites (**Fig. 3a, b**). The change in relative folding free energy (ΔΔG_fold_) served as a primary metric for evaluating the structural integrity and thermodynamic stability of the projected recombinants, a methodology widely validated for guiding the design of robust industrial enzymes. ^19^ As illustrated in Fig. 3a and b, we selected promising beneficial substitutions from the CASPE-T (S33T, S39T, D55N, Y57N, R249L, Y319L, Y320H, A372S, T413R, T513E, T526S) and CASPE-A (Q93E, S100N, M232Q, S234G, A304G, Q305N, Y341F, A357K, I382E, S419A, A429G, R436N) libraries with ΔΔG_fold_ < 0.36 kcal/mol for generating in silico comprehensive double-site recombinants. Notably, combinations involving T513E or T526S from the CASPE-T library paired with Q305N or Y341F from the CASPE-A library demonstrated the lowest ΔΔG_fold_, suggesting a favorable thermodynamic compatibility between these distant residues (**Fig. 3c**). Based on these computational insights, 10 double-site recombinants with the minimum predicted ΔΔG_fold_ were prioritized for experimental characterization.

**Fig. 3.**
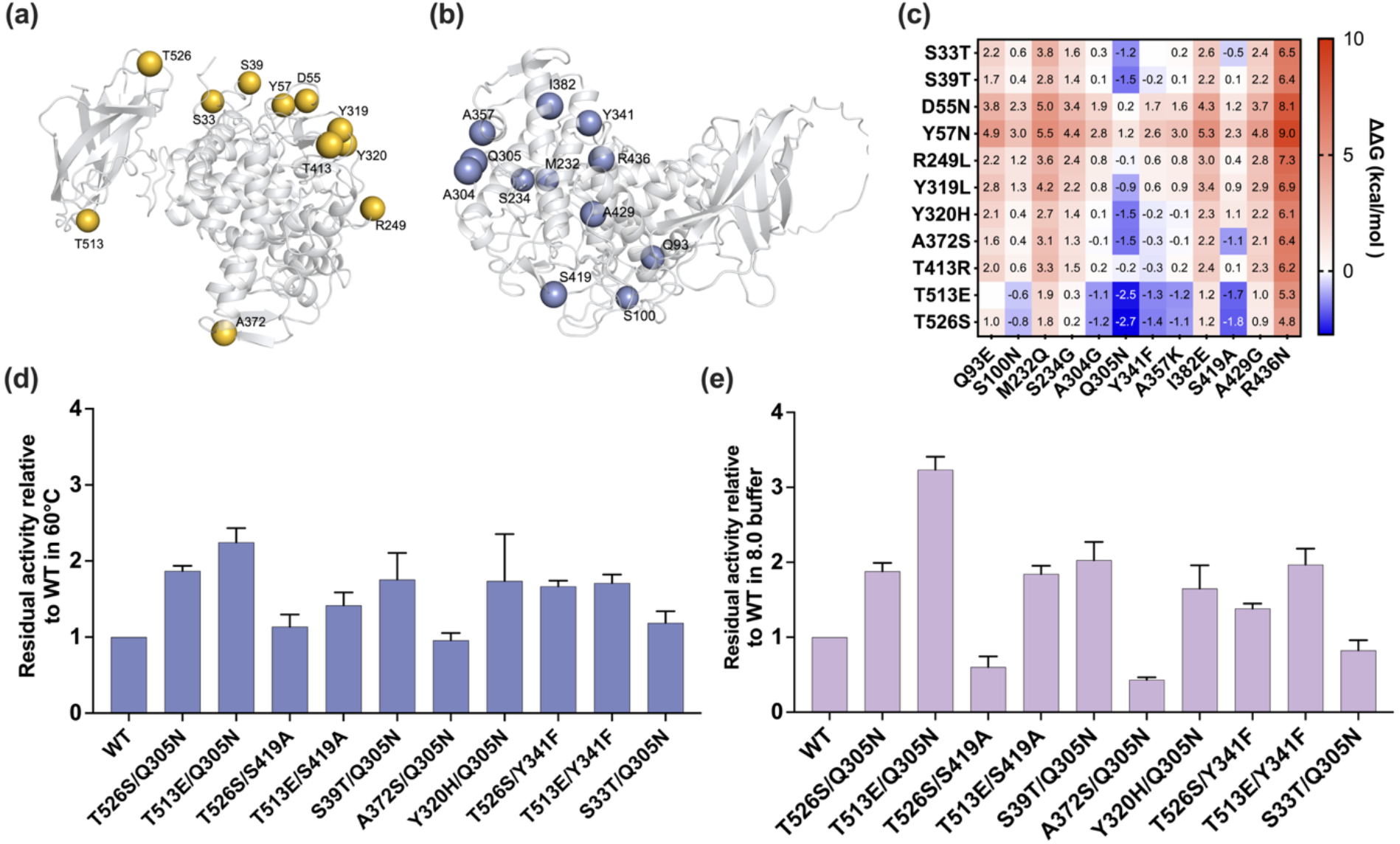
Identification and combinatorial recombination of beneficial mutations for dual-trait improvement. (a, b) Distribution of beneficial mutation sites identified by the thermostability-oriented (a) and alkaline stability-oriented (b) CASPE models, mapped onto the PoGA structure. (c) Prediction of changes in ΔΔG_fold_ for double-site recombinants using FoldX. The heatmap illustrates the thermodynamic stability effects of combining top-performing mutations from both models. (d, e) Experimental validation of selected recombinants. The residual activity of mutants relative to the WT after incubation at 60 °C (d) or in pH 8.0 buffer (e), respectively.

The results confirmed a strong correlation between the predicted ΔΔG_fold_ and the experimental robustness of the variants. The recombination strategy not only improved structural stability but also significantly enhanced catalytic activity. Among ten recombinants, nine of them exhibited thermal tolerance exceeding 1.1-fold that of the WT. The variants T526S/Q305N and T513E/Q305N performed exceptionally well, achieving 1.87-fold and 2.24-fold increases in activity compared to the WT, respectively (**Fig. 3d**). In addition, several other recombinants, including T526S/Y341F, T513E/Y341F, Y320H/Q305N, and S39T/Q305N, also maintained approximately 1.7-1.8-fold higher residual activity than the WT, demonstrating the broad effectiveness of the recombination strategy. Alkaline tolerance assessment further confirmed the effectiveness of this recombination strategy in adapting the enzyme to extreme pH environments (**Fig. 3e**). The optimal variant T513E/Q305N achieved a 3.24-fold increase in residual activity at pH 8.0, representing a significant enhancement over its single-point parents, T513E (~1.35-fold) and Q305N (~1.4-fold). Similarly, the recombinant S39T/Q305N exhibited a 2.03-fold increase in alkaline resistance, effectively surpassing the performance of S39T (~1.1-fold). Other recombinants, such as T513E/Y341F (1.97-fold) and T526S/Q305N (1.88-fold), also demonstrated superior pH stability compared to their isolated counterparts, successfully integrating the high alkaline tolerance of Y341F (~2.3-fold) into a more robust molecular framework.

Notably, while the majority of single point mutations maintained an initial catalytic activity comparable to the WT (**Fig. S1**), the recombinants exhibited a distinct enhancement in their intrinsic activity. Specifically, the optimal variant T513E/Q305N demonstrated an impressive 2.21-fold increase in specific activity compared to WT (**Fig. S2**). This observation indicates that the recombination strategy not only successfully integrated the stability traits but also synergistically boosted the baseline catalytic efficiency. These findings demonstrate that combining CASPE guided site identification with ΔΔG_fold_ based recombination effectively captures synergistic interactions among key residues, yielding variants with integrated improvements in activity and environmental tolerance.

### Thermostability and pH tolerance profile of the identified PoGA recombinants

To rigorously evaluate the industrial potential and structural robustness of the engineered PoGA, three PoGA recombinants (T513E/Q305N, T526S/Q305N, and T513E/Y341F) were subjected to thermostability and alkaline tolerance profiling. The recombinants profoundly altered the pH and temperature dependencies of PoGA. The pH activity profiles revealed a distinct alkaline shift, moving the optimal pH from 4.5 for the WT to a mildly acidic range of 5.0 to 5.5 for T513E/Q305N and T526S/Q305N, respectively (**Fig. 4a-c**). This upward adaptation facilitates seamless integration into existing industrial protocols, effectively bypassing the need for chemical pH adjustments during sequential starch processing.^24^ While the native enzyme exhibited a strict temperature optimum at 60 °C, the engineered recombinants, especially T513E/Y341F, displayed a broadened thermal activation peak extending to 65 °C. Furthermore, these variants maintained substantial catalytic activity at 75 °C, a stringent condition under which the native enzyme suffers rapid and complete inactivation (**Fig. 4d-f**).

**Fig. 4.**
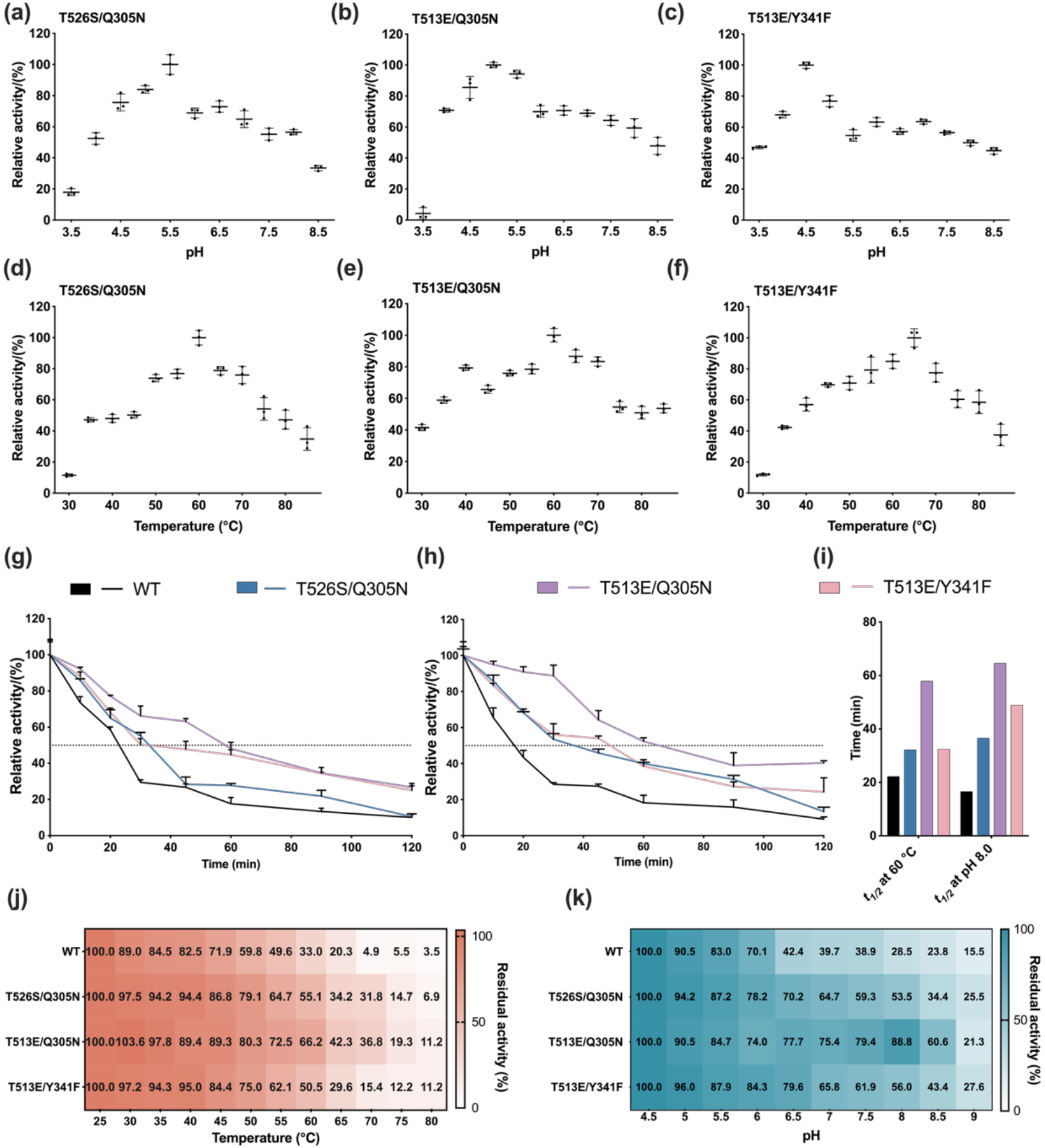
Comprehensive biochemical characterization and stability profiling of the optimal PoGA recombinants. pH activity profiles of the engineered variants: T526S/Q305N (a), T513E/Q305N (b), and T513E/Y341F (c). The optimal pH for each recombinant was determined by measuring catalytic activity across a pH range of 3.5 to 8.5. Temperature activity profiles of the engineered variants: T526S/Q305N (d), T513E/Q305N (e), and T513E/Y341F (f). The optimal reaction temperature was identified by assaying activity from 30 °C to 85 °C. The curves represent the thermal half life during incubation at 60 °C (g) and the alkaline half life during incubation at pH 8.0 (h) over time. The horizontal dashed lines indicate 50% residual activity. (i) Comparison of the *t*_*1/2*_ at 60 °C and pH 8.0 for the WT and the engineered variants. (j and k) Heatmap analysis of the thermostability (j) and pH stability (k) for the WT and the three optimal recombinants.

Kinetic stability assessments revealed a dramatic extension in resistance to unfolding under thermal and alkaline stress compared to the WT. At 60 °C, the WT exhibited a thermal half-life (*t*_*1/2*_) of only 22.29 min. In contrast, the variant T513E/Q305N achieved a remarkable 57.94 min, representing a 2.6-fold increase. The variants T513E/Y341F and T526S/Q305N also displayed significant improvements, reaching 32.46 min and 32.21 min, respectively. Under alkaline conditions at pH 8.0, the WT proved highly unstable with a *t*_*1/2*_ of only 16.6 min, whereas the engineered variants demonstrated superior resilience. Notably, the T513E/Q305N variant extended its alkaline half-life to 64.71 min, marking a nearly 4-fold improvement over the WT. The recombinant T513E/Y341F also exhibited substantial pH stability with a *t*_*1/2*_ of 48.92 min, while T526S/Q305N maintained a significant t_1/2_ of 36.64 min (**Fig. 4i-k**).

Condition stability profiles further corroborated this enhanced structural robustness by measuring residual activity following a 30 minute incubation across continuous temperature and pH gradients (**Fig. 4j, k**). The WT exhibited severe vulnerability to environmental stress, whereas the recombinant variants demonstrated profound resistance to thermal and alkaline unfolding. For instance, following incubation at 60 °C, T513E/Q305N preserved 66.2% residual activity, a 2.0-fold improvement over the WT. This thermal superiority became highly pronounced at 70°C, where the WT was nearly inactivated with only 4.9% activity remaining, while T513E/Q305N maintained 36.8% activity, achieving a remarkable 7.5-fold enhancement. Parallel quantitative improvements emerged in alkaline tolerance. After incubation at pH 6.5, T513E/Q305N retained 77.7% residual activity, equating to a 1.8-fold increase compared to the native enzyme. This structural resilience extended deeper into alkaline conditions. At pH 8.5, where WT activity was severely compromised to 23.8%, T513E/Q305N and T513E/Y341F maintained 60.6% and 43.4% residual activity, corresponding to 2.5-fold and 1.8-fold enhancements, respectively. These findings indicate that the combinatorial approach significantly enhanced the structural resilience of the enzyme, bringing its performance closer to the stringent demands of industrial starch processing.

### Molecular understanding of the superior performance

MD simulations were employed to systematically investigate the mechanisms underlying the superior thermostability and alkaline resistance of three identified PoGA recombinants (T526S/Q305N, T513E/Q305N, and T513E/Y341F). PoGA WT was also included for comparison. Dynamic structural analysis reveals the primary conformational transitions occurring over time. As illustrated in **Fig. 5a**, under standard conditions, the root-mean-square deviation (RMSD) values of the three variants (~6.2–8.0 Å) remained significantly lower than those of the WT (~12.0–12.7 Å), a trend that persisted at both pH 4.5 and pH 8 (**Fig. S3**). Notably, the WT exhibited an abrupt decrease in RMSD, signaling a sharp structural shift under thermal stress at 60 °C, whereas the recombinants did not. This enhanced global stability is further confirmed by the analysis of internal hydrogen-bond networks (**Fig. 5b, S4, and S6**). Specifically, the T513E/Q305N variant exhibited the highest number of high-occupancy hydrogen-bonds (occupancy > 95%) across all simulated conditions. This observation indicates that the recombinants significantly stabilize the essential hydrogen-bonding scaffold, more effectively anchoring secondary structural elements and increasing overall structural rigidity ^42, 43^. Consistent with these findings, the free energy landscape (**Fig. 5g**) for T513E/Q305N shows a deeper and more concentrated minimum energy basin, indicating a more stable conformational ground state ^44^.

**Fig. 5.**
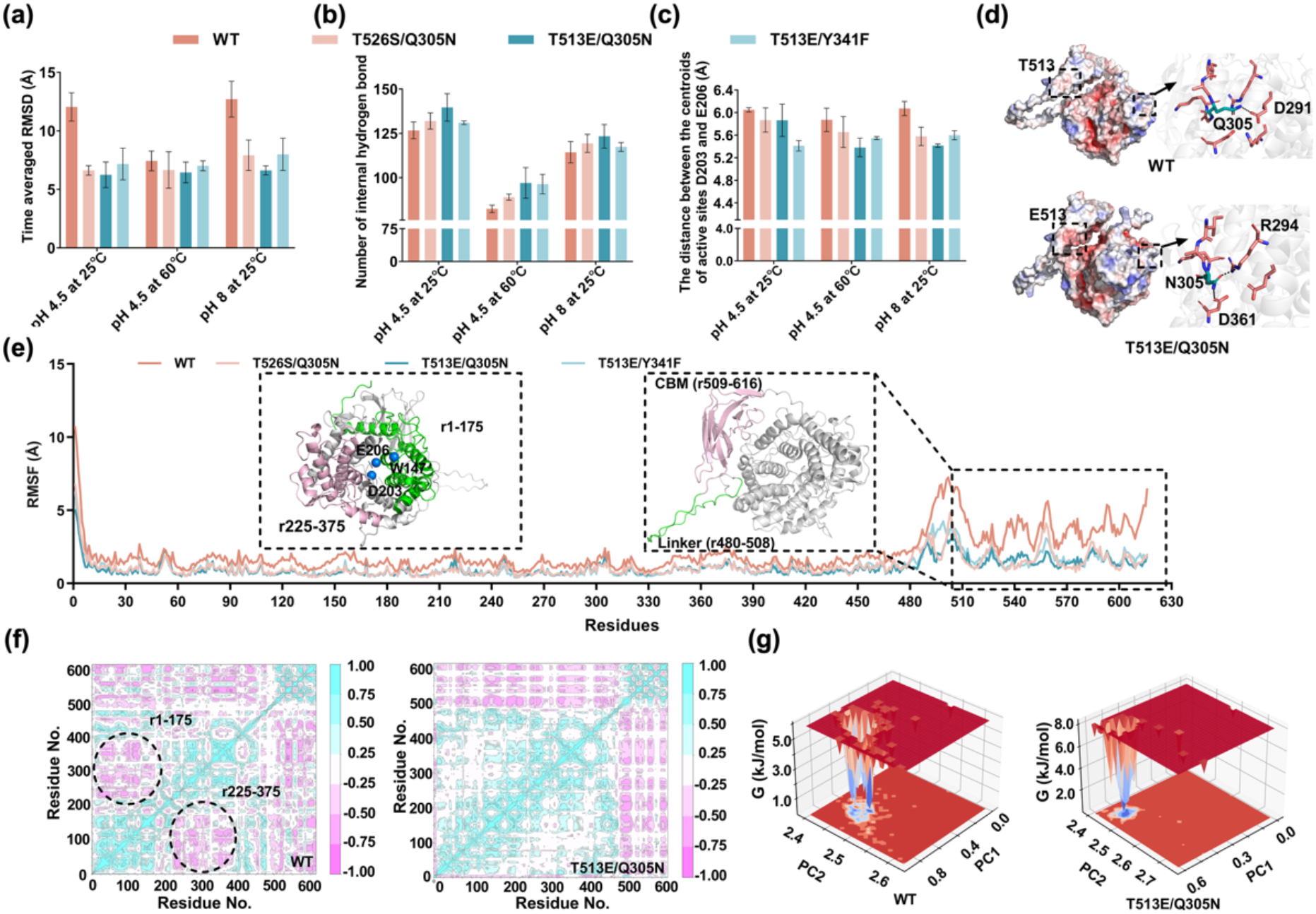
MD simulations elucidate the mechanism of enhanced thermostability and alkaline resistance in recombinant variants through structural insights. (a) Time averaged root-mean-square deviation (RMSD) of WT and the recombinant variants T526S/Q305N, T513E/Q305N, and T513E/Y341F backbone (Cα, N, and carbonyl C) with respect to the initial structure in different temperature and pH conditions during the last 40 ns of MD simulation. (b) The number of internal hydrogen bonds with >95% occupancy within enzymes in different temperature and pH conditions during the last 40 ns of MD simulation. (c) Average distance between the centers of mass of the active sites D203 and E206 in different temperature and pH conditions during the last 40 ns of MD simulation. (d) The electrostatic potential energy surfaces of WT and T513E/Q305N, with red indicating a negative charge, blue indicating a positive charge, and white indicating a net charge of 0. Two mutated sites were boxed, and the hydrogen bonding network of residues surrounding Q305N was further illustrated. Residues interacting with Q305N via hydrogen bonds were labeled, with hydrogen bonds depicted as black dashed lines. (e) RMSF of WT and the recombinant variants T526S/Q305N, T513E/Q305N, and T513E/Y341F residues at 25 °C, pH=8 during the last 40 ns of MD simulation. Highlighted were the residue regions with larger fluctuations and their structural diagrams, with the linker and CBM domain shown in different colors. Two helix clusters within the main domain are highlighted, along with the enclosed catalytic triad (W147, D203, and E206), with their Cα atoms shown as blue spheres. (f) Dynamical Cross-Correlation Matrix (DCCM) of WT and T513E/Q305N under optimal conditions, with differential regions circled. (g) Free energy landscape (FEL) analysis of WT and T513E/Q305N under pH 4.5 at 25 °C. PC1 represents the RMSD value; PC2 represents the R_g_ value, and the 3D FEL of the enzyme was represented by using PC1 and PC2 of the enzyme as the reaction coordinates. Data plotted from the average of three independent MD runs.

The transition from global stability to localized resilience is evident in the domain-specific fluctuation profiles. Root-mean-square fluctuation (RMSF) analysis (**Fig. 5e and S5**) shows that while the catalytic domain (residues 1–479) remains rigid in all systems, the linker (residues 480–508) and the CBM (residues 509– 616) serve as the key sources of conformational instability of the WT. At pH 8.0 and 60°C, the β-sheet-rich CBM in the WT undergoes a collapse of its inter-strand hydrogen-bonding network due to thermal agitation and electrostatic repulsion. In contrast, the variants maintain low-fluctuation profiles across these regions. This stabilization is further explained by dynamic cross-correlation matrix (DCCM) analysis (**Fig. 5f**). In the WT, two helical segments within the catalytic domain (residues 1–175 and 225–375) exhibit anticorrelated motions, suggesting internal incoordination. Conversely, the T513E/Q305N variant shifts toward strong positive correlation, indicating that distal mutations induce long-range allosteric effects that lock the catalytic domain into a more cooperative and robust conformation (**Fig. 5e**). This enhanced dynamic coordination is directly reflected in the structural integrity of the active site.

Center-of-mass distance analysis between the catalytic residues D203 and E206 (**Fig. 5c and S8**) reveals that the distances in the three variants are consistently shorter than those in the WT. Under harsh environmental conditions, the T513E/Q305N variant maintains the most compact catalytic pocket, exhibiting the minimum distance observed across all systems. This suggests that the synergistic effects of the substitutions effectively prevent the unfavorable dynamics within the active site that occur in the WT due to thermal or pH-induced fluctuations. By maintaining an optimized proximity between the nucleophile and the proton donor, the variants—particularly T513E/Q305N—ensure efficient proton transfer, thereby providing a robust structural basis for their superior catalytic efficiency under stress ^45, 46^. More description about the corresponding surface electrostatic interactions was described in Supporting Information (**Text S1, Fig. 5d, and Fig. S7-S9, S12**).

In summary, the enhanced performance of the engineered PoGA variants originates from a cascade of structural stabilization events spanning multiple spatial scales. Reinforcement of the internal hydrogen-bonding network establishes a more rigid global scaffold, which suppresses conformational fluctuations in the linker and CBM while simultaneously enhancing cooperative motions within the catalytic domain. These collective dynamic changes ultimately preserve the integrity and compactness of the catalytic pocket, enabling efficient proton transfer and sustained catalytic function under elevated temperature and alkaline conditions. Therefore, the improved thermostability and alkaline resistance arise not from a single local interaction, but from synergistic multiscale reorganization of the protein structural dynamics.

### Mass balance of saccharification performance

To evaluate the industrial applicability of the engineered glucoamylase, three starch liquefaction– saccharification process configurations were designed. To enable a direct comparison of glucose production among the three configurations, the experimentally determined glucose yields were further used to establish mass balances on the basis of 1 t of dry corn flour. In all configurations, starch liquefaction was performed under typical industrial conditions (95 °C and pH 6.5). After liquefaction, the hot slurry was first cooled from 95 °C to 60 °C by heat exchange with recovered process water (RPW). The heated RPW was subsequently reused as slurry preparation water for the next liquefaction batch, thereby recovering sensible heat and reducing the thermal energy required for slurry heating in the subsequent liquefaction process. Following this common heat recovery step, three saccharification configurations were evaluated: Process I, in which the slurry was further cooled from 60 °C to 30 °C using recirculating cooling water (RCW) and adjusted from pH 6.5 to 4.5; Process II, in which saccharification was performed at 60 °C and pH 4.5 after acidification; and Process III, in which saccharification was directly performed at 60 °C and pH 6.5 without acidification.

The resulting mass balances highlighted distinct performances of WT and T513E/Q305N under the three process configurations. Under Process I (30 °C and pH 4.5, **Fig. 6a**), WT produced 158.4 g/L glucose, corresponding to 527.9 kg of glucose from 1 t of dry corn flour. In contrast, T513E/Q305N increased the glucose concentration to 199.3 g/L, 25.8% higher than WT, while achieving a starch conversion of 80.8%, compared with 64.2% for WT. Under Process II (60 °C and pH 4.5, **Fig. 6b**), the glucose concentration of T513E/Q305N further increased to 227.5 g/L, compared with 192.5 g/L for WT, while the starch conversion reached 92.2%, compared with 78.0% for WT. These results demonstrate that T513E/Q305N maintains high saccharification efficiency under both low- and high-temperature conditions, highlighting its potential for low-temperature saccharification compatible with simultaneous saccharification and fermentation as well as high-temperature saccharification applicable to separate hydrolysis processes ^47, 48^.

**Fig. 6.**
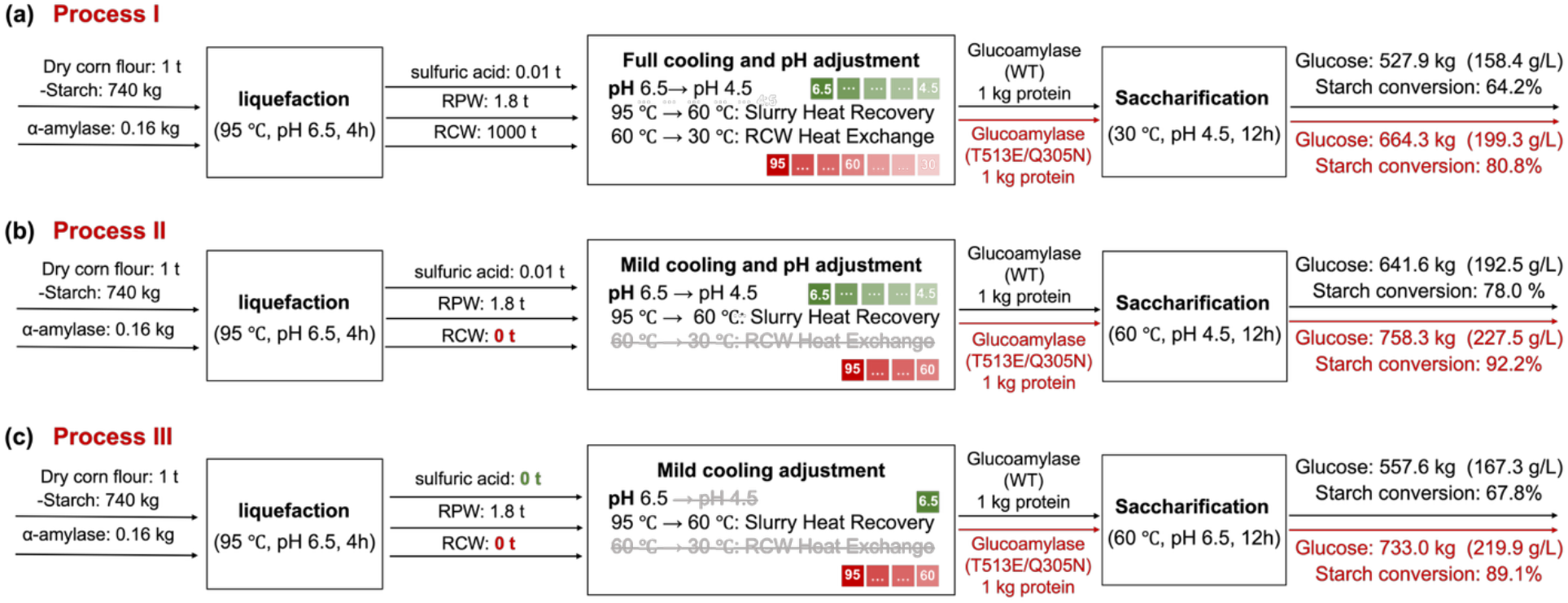
Mass balance of glucose production from 1 metric ton of dry corn powder under three starch liquefaction and saccharification process configurations using the PoGA WT or the T513E/Q305N variant. Industrial process parameters were referenced from SDIC Biotech Tieling. In all cases, starch liquefaction was performed at 95 °C and pH 6.5, whereas saccharification was evaluated under varying operational conditions. (a) Process I requires initially cooling the liquefied slurry to 60 °C using recycled process water (RPW). The RPW is simultaneously used for the next liquefaction mashing step to reduce subsequent energy consumption. The slurry is then further cooled to 30 °C using recirculating cooling water (RCW) and adjusted to pH 4.5 by adding acid prior to saccharification. (b) Process II uses RPW to reduce the temperature of the liquefied slurry to 60 °C, followed by acid addition to achieve a pH of 4.5 for saccharification. (c) Process III employs RPW-based heat recovery to cool the slurry to 60 °C and proceeds directly to saccharification without any acid addition or pH adjustment.

Given the excellent saccharification performance of T513E/Q305N under both Process I and Process II, we further evaluated whether saccharification could be performed directly at the native liquefaction pH without acidification (**Fig. 6c)**. Accordingly, Process III was conducted at 60 °C and pH 6.5 without sulfuric acid addition. Under these conditions, T513E/Q305N maintained a high glucose concentration of 219.9 g/L, which was 31.4% higher than that of WT (167.3 g/L), while achieving a starch conversion of 89.1% compared with 67.8% for WT. More importantly, eliminating sulfuric acid addition resulted in only a 3.1% reduction in starch conversion for T513E/Q305N, compared with a 10.2% reduction for WT relative to Process II. These results demonstrate that T513E/Q305N retains high saccharification efficiency at the native liquefaction pH, suggesting its potential to enable direct saccharification following liquefaction without acidification, thereby reducing chemical consumption and simplifying industrial starch processing. Collectively, these results demonstrate that simultaneous enhancement of thermostability and alkaline tolerance can be translated into tangible process-level benefits, providing a practical strategy for developing robust industrial glucoamylases with broad potential in starch processing, ethanol and glucose/fructose syrup production, as well as textile, detergent, and paper industries ^49^.

## Conclusion

In this study, we established an artificial intelligence-guided framework for simultaneously improving the thermostability and pH tolerance of PoGA. The property-specific CASPE-T and CASPE-A models identified largely distinct but partially coupled mutational landscapes, demonstrating that these two traits arise from overlapping yet non-identical structural determinants. Combining experimentally validated substitutions through folding-energy-guided recombination enabled favorable interactions between distant mutation sites and generated variants with integrated improvements in stability and catalytic activity. Among the engineered enzymes, T513E/Q305N showed the best overall performance, exhibiting a 2.21-fold increase in specific activity, a 2.6-fold extension of its half-life at 60 °C, and an approximately 3.9-fold extension of its half-life at pH 8.0 relative to the WT.

MD simulations indicated that these improvements resulted from multiscale reorganization of protein dynamics rather than a single localized interaction. Reinforced hydrogen-bonding networks stabilized the global scaffold, reduced fluctuations within the linker and carbohydrate-binding module, promoted cooperative motions across the catalytic domain, and preserved the compact geometry of the catalytic residues under thermal and alkaline stress. Importantly, these molecular improvements translated into enhanced process performance. At 60 °C and the native liquefaction pH of 6.5, T513E/Q305N achieved 89.1% starch conversion and produced 219.9 g/L glucose without acidification. The engineered enzyme could therefore simplify the transition from liquefaction to saccharification and reduce chemical consumption under the tested conditions. Overall, this work provides both a promising glucoamylase candidate for starch processing and a transferable strategy for multi-objective enzyme engineering through the integration of property-specific prediction, thermodynamic filtering, experimental validation, and mechanistic simulation. Further pilot-scale evaluation, long-term operational testing, and technoeconomic analysis will help establish its industrial applicability.

## Acknowledgments

This work was supported by the National Key Research and Development Program of China (2024YFA0917800), the Coal-Major Project (Grant No. 2025ZD1701600), Basic Research Program of Jiangsu Province (Grant NO. BK20253050, BK20250141), Qinglan Project of Jiangsu Province of China, Jiangsu Basic Research Center for Synthetic Biology (BK20233003), State Key Laboratory of Microbial Technology Open Projects Fund (Grant No. M2025-17)).

## Declaration

The authors declare that they have no conflict of interest.

