## Supplementary Materials for "AI-Guided Multi-Objective Engineering of Glucoamylase Enables Acidification-Free Starch Saccharification"

[c] Q. Deng
School of Environmental and Biological Engineering
Nanjing University of Science and Technology
Xiaolingwei Street, Nanjing 210094, China

[d] Dr. G. Liu
State Key Laboratory of Microbial Technology
Shandong University
No. 72 Binhai Road, Qingdao, 266237, China

[e] F. Shi, L. Deng
SDIC Bioenergy Tieling CO., LTD.
Tieling, 112700, China


#These authors contributed equally

### Materials and Methods

#### Materials

The restriction enzyme *Dpn*I and PrimeSTAR High-Fidelity DNA Polymerase were purchased from Takara (Dalian, China). Soluble starch was obtained from Merck Ltd. (Beijing, China), and the DNS reagent was purchased from Beijing Solarbio Science & Technology Co., Ltd. (Beijing, China). All other chemicals were of analytical grade and were acquired from Shanghai Macklin Biochemical Co., Ltd. (Shanghai, China).

#### Strain and plasmid

The gene encoding GA from *Penicillium oxalicum* was kindly provided by Prof. Guodong Liu in Shandong University (Shandong, China). The in-house plasmid vector pPICZαA-GAP was used for protein expression [1]. *Escherichia coli* DH5α was used for plasmid propagation and mutant construction, while *Pichia pastoris* X-33 served as the host strain for the expression of both wild type (WT) and mutant enzymes.

#### Protein expression procedure in 96-well MTP

High-throughput expression of PoGA in *P. pastoris* X33 was performed using 96-well flat-bottom microplates. The strains were cultivated in YPD broth (1% yeast extract, 2% peptone, and 2% glucose) containing 100 μg/mL Zeocin. For fermentation, 5 μL of the pre-culture (grown for 48 h at 30 °C, 900 rpm) was inoculated into 160 μL of fresh YPD medium. The main culture was incubated for 96 h under the same conditions (25 °C, 900 rpm). Finally, the supernatant was collected by centrifugation at 2,000 × g and 4 °C for 20 min and transferred to clean plates for further assays.

#### Stability assays and activity measurement PoGA

PoGA activity was assayed in 96-well PCR plates using a modified DNS method according to a previous study [2]. A reaction mixture containing 10 μL of enzyme and 90 μL of 2% (w/v) soluble starch was incubated for 5 min. Following the addition of 100 μL of DNS reagent, the mixture was heated at 95 °C for 15 min and cooled at 10 °C for 10 min. The absorbance of a 100 μL final aliquot was then recorded at 540 nm.

To determine the optimal pH, enzyme activity was measured in various buffers covering a broad pH range: 0.1 M sodium acetate (pH 3.5 to 5.5), 0.1 M sodium phosphate (pH 6.0 to 8.0), and 0.1 M Tris HCl (pH 8.5 to 9.5). The relative activity was defined as the ratio of the activity at a specific pH to the maximum activity observed at the optimal pH (set as 100%). To determine the optimal temperature, the enzyme assay was performed at temperatures ranging from 25 to 85 °C at the optimal pH. The relative activity was calculated by normalizing the activity at a specific temperature to the maximum activity at the optimal temperature (set as 100%).

To determine pH stability, the enzyme solution was diluted 5-fold in the aforementioned buffers and incubated at room temperature for 0.5 h with agitation at 900 rpm. Subsequently, a 10 μL aliquot of the treated mixture was used for the standard activity assay. Residual activity was calculated by normalizing the activity of the treated samples to that of the control sample incubated at pH 4.5 (defined as 100%). For thermostability analysis, the enzyme was diluted 5-fold in 0.1 M sodium acetate buffer (pH 4.5) and incubated at designated temperatures ranging from 25 to 80 °C for 0.5 h with agitation at 900 rpm. A 10 μL aliquot was then subjected to the activity assay. Residual activity was assessed by normalizing the activity of the heat treated samples to that of the untreated control incubated at 25 °C (defined as 100%).

**Mutant construction**

Site-directed mutagenesis (SDM) was performed on the PoGA WT plasmid by PCR, according to the QuikChange mutagenesis method [3]. Primers are listed in the Supplementary Information.

**Corn liquefaction and saccharification**

A corn slurry was prepared at a concentration of 30% (w/v) based on the absolute dry weight of the corn flour. Commercial α-amylase was added to the mixture, and the initial pH was adjusted to 6.5. The liquefaction process was then conducted in a water bath at 95 °C for 4 h.

Following liquefaction, the slurry was divided into three groups for saccharification at (i) 30 °C and pH 4.5, (ii) 60 °C and pH 4.5, and (iii) 60 °C and pH 6.5, respectively. WT and the engineered variant T513E/Q305N were evaluated under all three process conditions. The amount of WT or engineered glucoamylase added was normalized according to enzymatic activity to match the activity provided by the commercial glucoamylase at a dosage of 0.1% (w/w, dry corn flour basis). All saccharification reactions were performed for 12 h with constant agitation at 250 rpm. The experimentally determined glucose production of WT and T513E/Q305N under the three saccharification conditions was used to construct the corresponding mass balances for glucose production using 1 t of dry corn flour as the feedstock.

**Molecular dynamics simulations of the enzyme under different temperature and pH conditions**

Molecular dynamics (MD) simulations were performed using GROMACS 2023.3 [4] with Amber 99sb-ildn force fields [5]. A total of 36 molecular dynamics simulations were performed for four PoGA variants under two temperature and pH conditions. To simulate two distinct pH environments (pH 4.5 and pH 8.0), protonation state assignments were performed for PoGA WT and three variants PoGA T526S/Q305N, T513E/Q305N, and T513E/Y341F. The pKa values of all titratable residues in PoGA were determined using Propka 2.0 [6], and the corresponding protonation states were specified via the interactive options in GROMACS 2023.3 to achieve the desired pH conditions. The enzyme was first placed in a cubic box with a minimum distance of 12 Å from the edge of the box to the proteins, and then the box was filled with the SPCE water molecules [7]. To equilibrate the system, $Na^+$ and $Cl^-$ were added to achieve net charge neutralization [8, 9]. The canonical ensemble (NVT) utilized the v-rescale heat bath method [10], whereas the isothermal-isobaric ensemble (NPT) employed the Parrinello–Rahman pressure controller [11]. To avoid unfavorable interactions, energy minimization was performed using the steepest descent method prior to performing the MD simulation, followed by NVT and NPT equilibrations for 100 ps. Finally, the simulation was run at 100 ns, 298 K/333 K, 1 bar, and 2 fs. To account for stochastic variations arising from a single simulation, three independent molecular dynamics simulations were performed using different initial atomic velocity distributions. During the simulations, energies, coordinates, and velocities were recorded at 0.5-ns intervals, and all analyses were calculated using the GROMACS simulation package tool. Finally, the trajectories were visualized and analyzed using PyMOL 3.1.3.

**Text S1** Analysis of Protein Surface Electrostatic Interactions

The observed structural robustness is fundamentally rooted in the specific physicochemical alterations at the residue level [12, 13]. The T513E/Q305N variant represents the optimal engineering strategy, combining mutations in both the catalytic domain and the CBM. In the catalytic domain, the Q305N substitution facilitates more precise hydrogen bonding due to its compact side chain (**Fig. 5d**). Simultaneously, the T513E substitution in the CBM introduced a negatively charged glutamic acid that strengthens surface electrostatic interactions (**Fig. 5d**). The synergy between these two sites might forming the molecular locker that rigidifies the linker-CBM interface. Other modifications contribute through distinct mechanisms: Y341F (catalytic domain) reduces conformational degrees of freedom by removing a hydroxyl group **(Fig. S12)**, while T526S (CBM region) alleviates steric hindrance to optimize local packing **(Fig. S12)**. Differences in hydration resulting from these recombinants are also reflected in **Figs. S7-S9**. Collectively, these strategic point mutations reconfigure the protein's microenvironment, driving the marked improvements in thermostability and alkaline resistance.

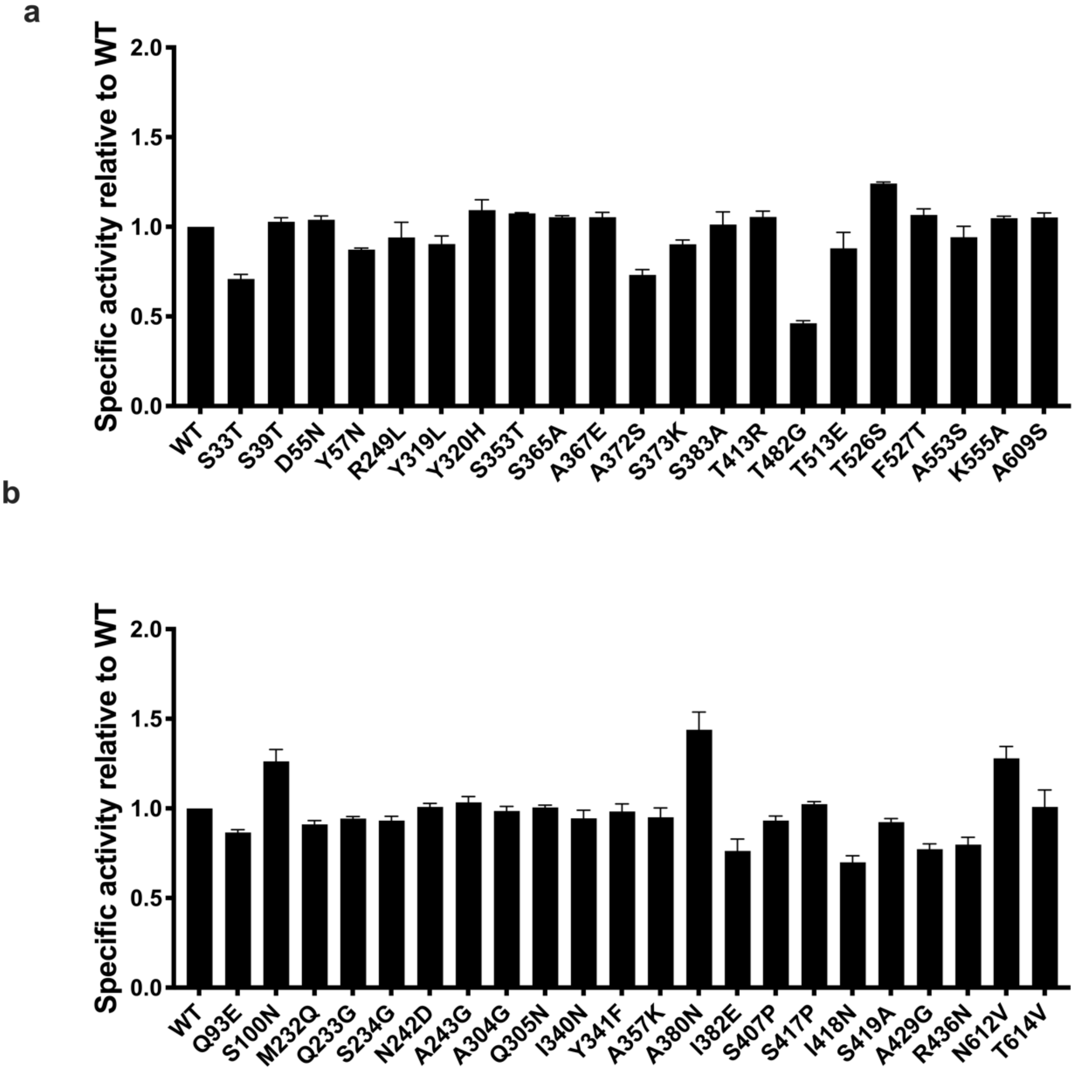

**Fig. S1** Specific activity of PoGA variants relative to the wild type (WT). (a) Mutants prioritized by the CASPET model for thermal stability. (b) Mutants prioritized by the CASPEA model for pH stability. Data for each variant are presented relative to the WT, which is normalized to 1.0. Error bars represent standard deviations from three independent experiments.

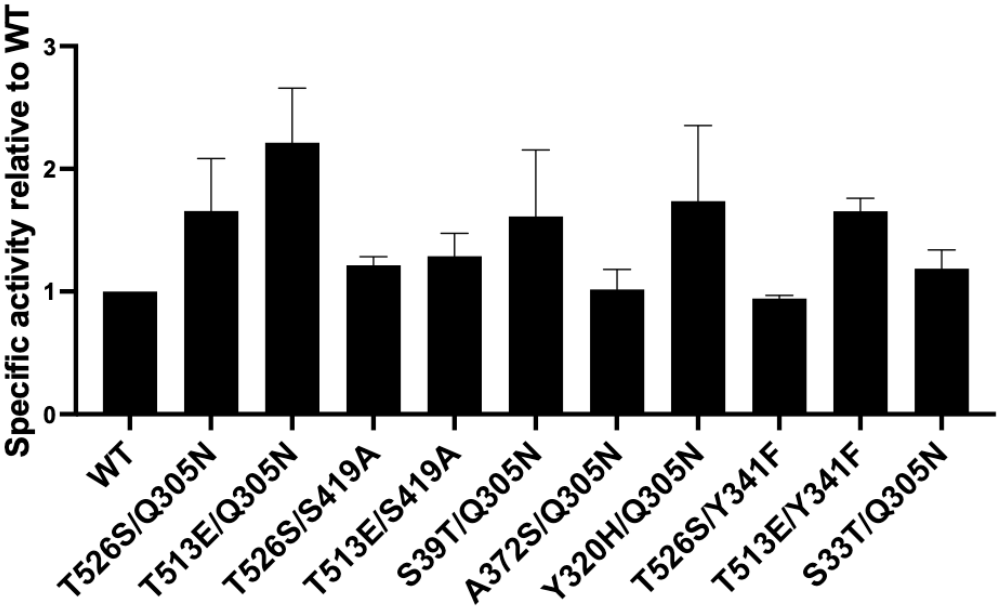

**Fig. S2** Specific activity of combinatorial PoGA variants relative to the WT. Error bars represent standard deviations from three independent experiments.

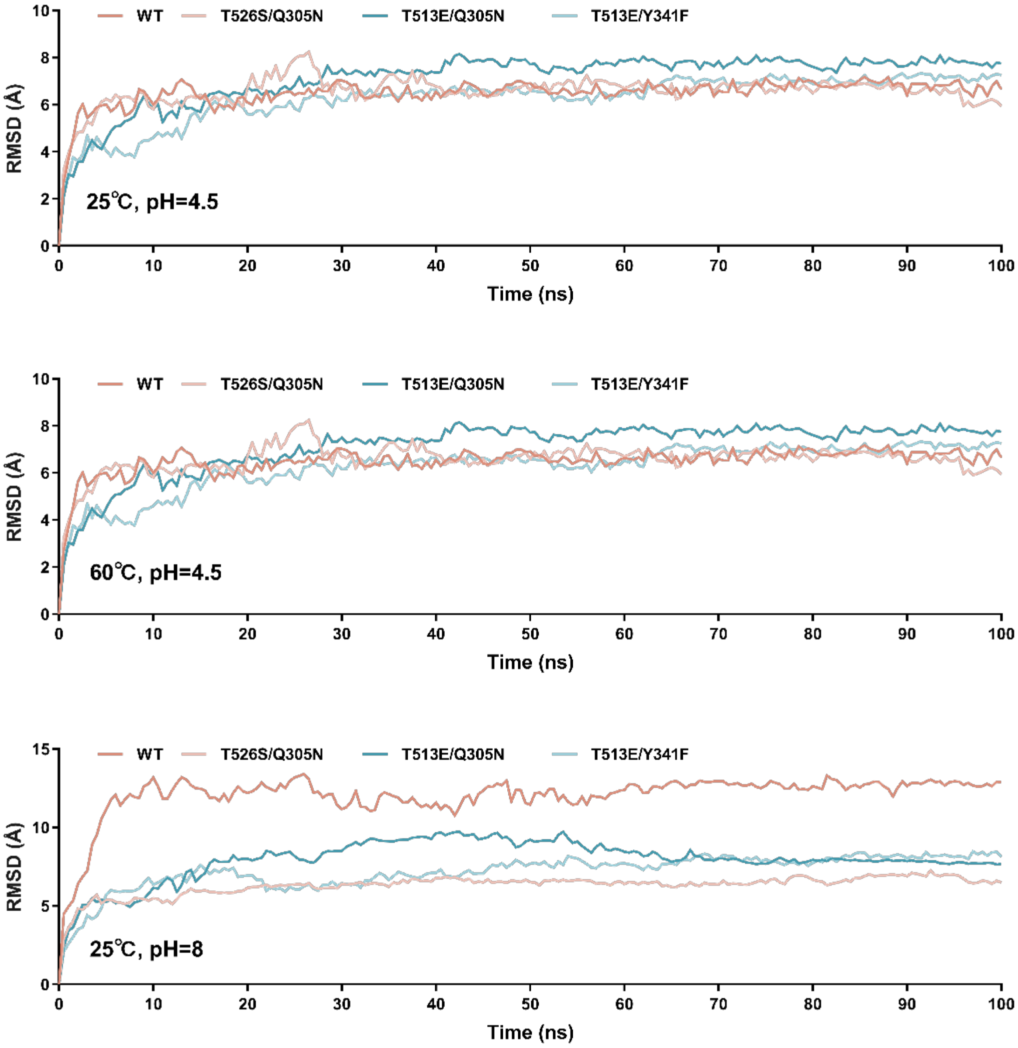

**Fig. S3** Root-mean-square deviation (RMSD) of the PoGA-WT, T526S/Q305N, T513E/Q305N, and T513E/Y341F backbone with respect to the initial structure in three environments during the last 40 ns of MD simulation, determined from three independent MD simulations.

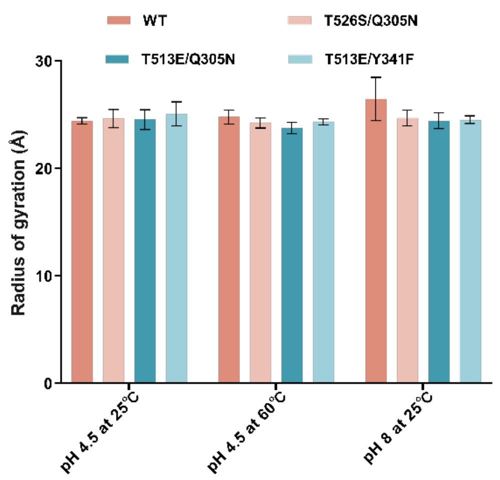

**Fig. S4** The radius of gyration (Rg) of the PoGA-WT, T526S/Q305N, T513E/Q305N, and T513E/Y341F with respect to the initial structure as a function of time in three environments during the last 40 ns of MD simulation. Error bars correspond to the standard deviation (s.d.) of three independent MD runs (n=3).

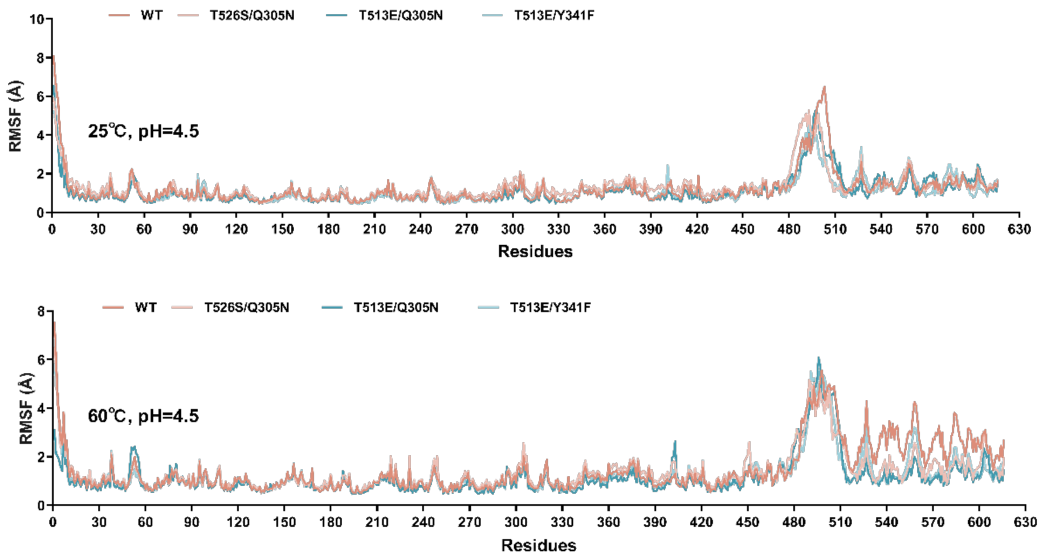

**Fig. S5** Root mean square fluctuation (RMSF) of the PoGA-WT, T526S/Q305N, T513E/Q305N, and T513E/Y341F backbone with respect to the initial structure in two environments during the last 40 ns of MD simulation, determined from three independent MD simulations.

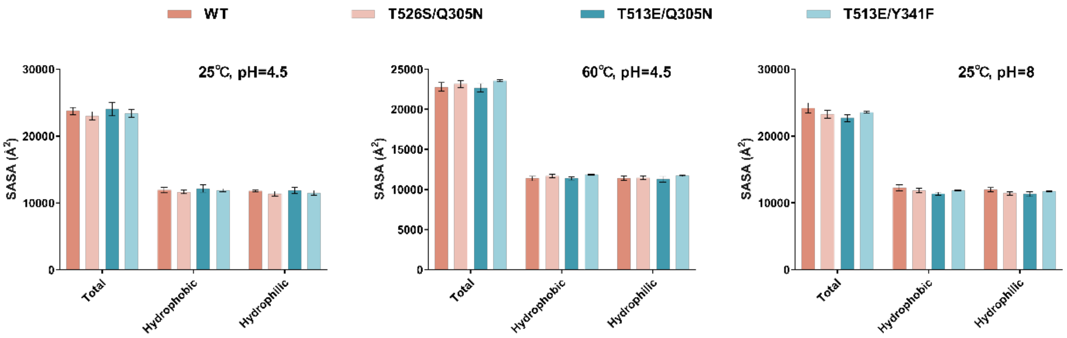

**Fig. S6** The time-averaged total solvent accessible surface area (SASA), the time-averaged hydrophobic SASA, and hydrophilic SASA of the PoGA-WT, T526S/Q305N, T513E/Q305N, and T513E/Y341F in three environments during the last 40 ns of MD simulation. Error bars correspond to the standard deviation (s.d.) of three independent MD runs (n=3).

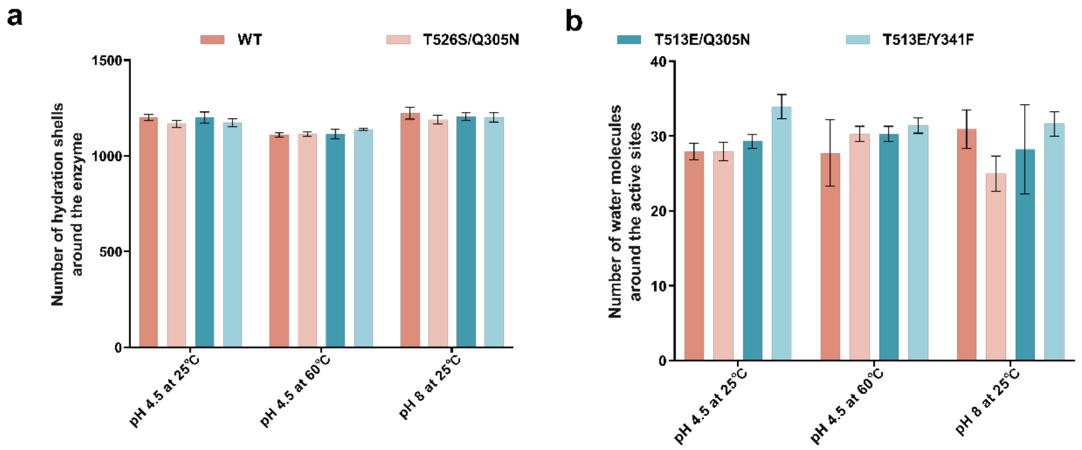

**Fig. S7** Hydration of enzymes in MD simulations. (**a**) The number of hydration shell of the PoGA-WT, T526S/Q305N, T513E/Q305N, and T513E/Y341F in three environments during last 40 ns of MD simulation. The hydration shell was defined as water molecules whose oxygen atom was localized at a distance less than 3.5Å from any nonhydrogen atom of the enzyme. (**b**) Number of water molecules around the active site (D203) of the PoGA-WT, T526S/Q305N, T513E/Q305N, and T513E/Y341F in three environments during last 40 ns of MD simulation. The distance around D203 was delineated as 8.1 Å, which was predicted by the Depth Server (http://cospi.iiserpune.ac.in/depth).[14]

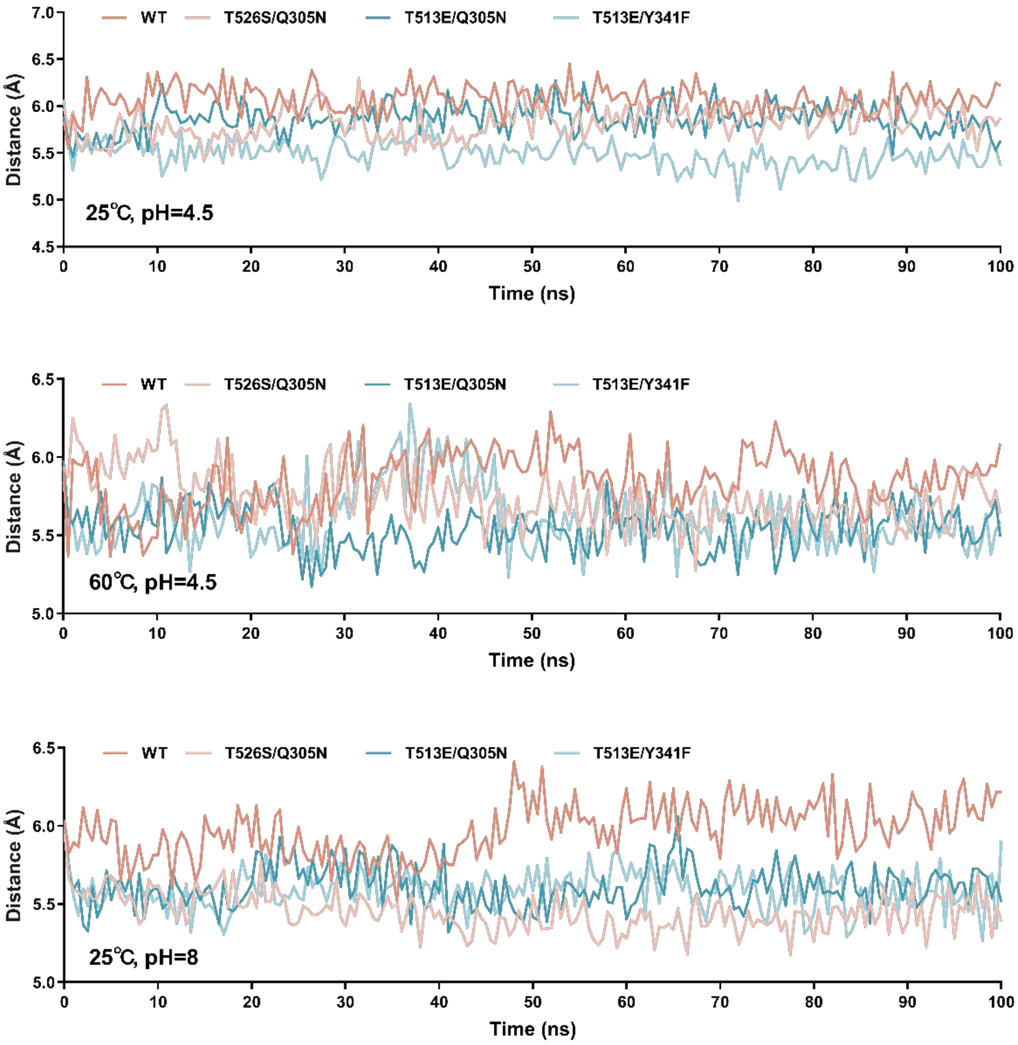

**Fig. S8** Distance between the centers of mass of the active sites D203 and E206 in three environments during the 100 ns of MD simulation, determined from three independent MD simulations.

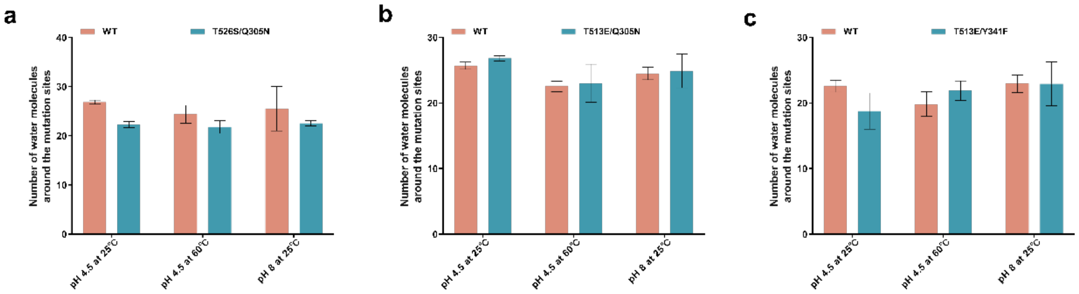

**Fig. S9** Number of water molecules around the mutation site (Q305N, Y341F, T513E, and T526S) of the PoGA-WT, T526S/Q305N, T513E/Q305N, and T513E/Y341F in three environments during last 40 ns of MD simulation. (a) WT and T526S/Q305N, (b) WT and T513E/Q305N, and (c) WT and T513E/Y341F. The number of water molecules was the sum of those at the two mutation sites. The distance around Q305N, Y341F, T513E, and T526S was delineated as 3.7 Å, 4.3 Å, 3.5 Å, and 3.7 Å, which was predicted by the Depth Server (http://cospi.iiserpune.ac.in/depth). Error bars correspond to the standard deviation (s.d.) of three independent MD runs (n=3).

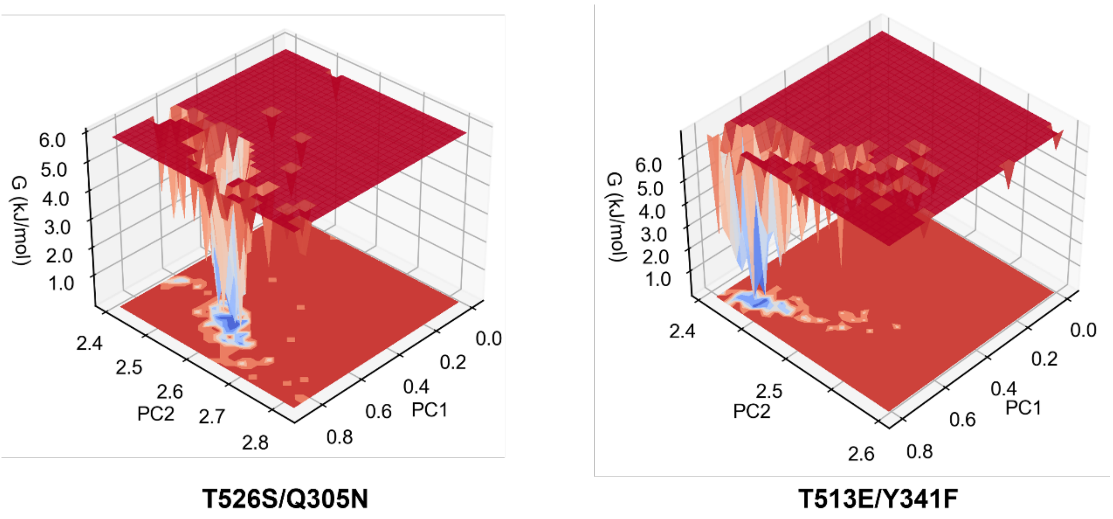

**Fig. S10** Free energy landscape (FEL) analysis of T526S/Q305N and T513E/Y341F under optimal conditions. PC1 represents RMSD value; PC2 represents Rg value, and the 3D FEL of the enzyme was represented by using PC1 and PC2 of the enzyme as the reaction coordinates.

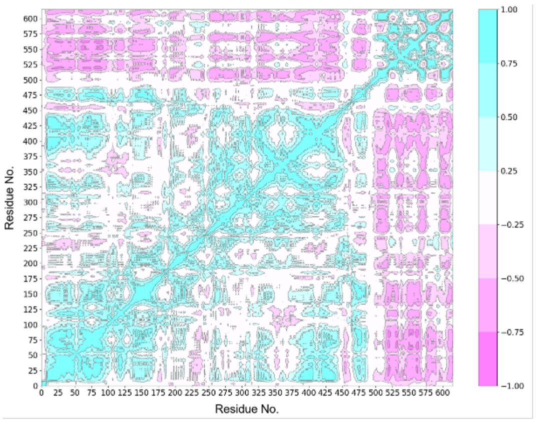

**T526S/Q305N**

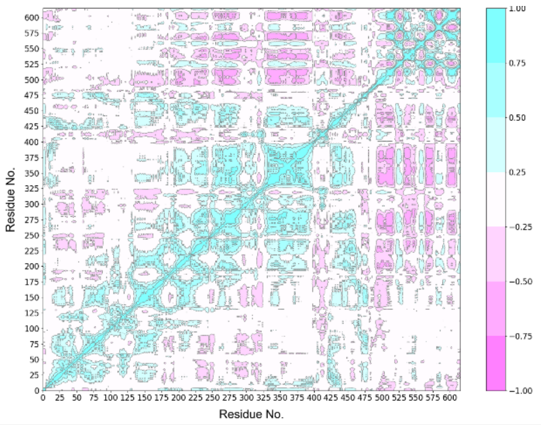

**T513E/Y341F**

**Fig. S11** Dynamical Cross-Correlation Matrix (DCCM) of T526S/Q305N and T513E/Y341F under optimal conditions.

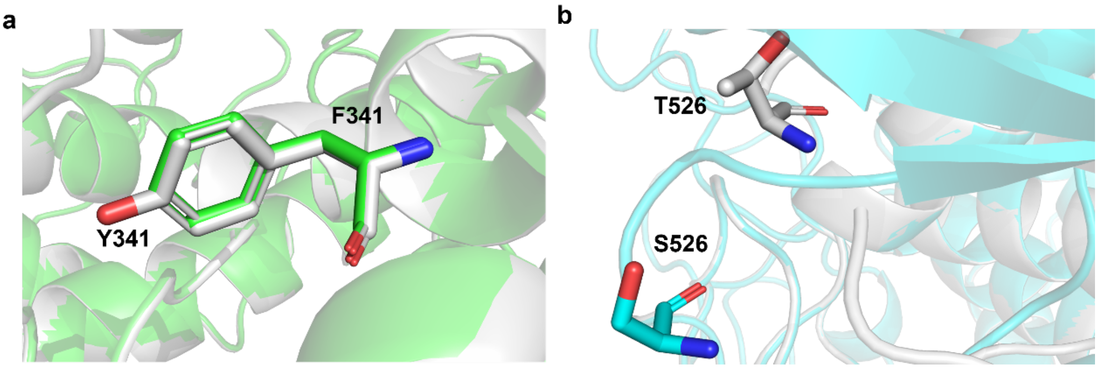

**Fig. S12** Superimposed structures of the last frames from MD simulations of the recombinant variant and WT, highlighting the changes at the mutation sites. WT was shown in gray, and the variant was shown in color. (a) WT and T513E/Y341F, (b) WT and T526S/Q305N. Data plotted from the average of three independent MD runs.

**Table S1.** Primers used for the construction of PoGA variants prioritized by the CASPET model.

| Name | 5' --> 3' sequence |
|---|---|
| S33T-F | ACATTGGCTCCaccGGTGCCTACTCCAAGAGTGC |
| S33T-R | gcACCggtGGAGCCAATGTTGGCCAGGATA |
| S39T-F | TGCCTACTCCAAGaccGCCGCCAGTGGTGCGGTCat |
| S39T-R | CGGCggtCTTGGAGTAGGCACCGCTGGAGCCAATG |
| D55N-F | CCAGCCCTaacTATTACTACACCTGGACCCGCG |
| D55N-R | GTAATAgttAGGGCTGGATGTGCTAGGGC |
| Y57N-F | CAGCCCTGACTATAACTACACCTGGACCCGCGACG |
| Y57N-R | GGGTCCAGGTGTAGTTATAGTCAGGGCTGgatgtgcta |
| R249L-F | GCcttTCCGGCAAGGACTCCAACACCA |
| R249L-R | AGTCCTTGCCGGAaagGCCACCGCCAGTGTTG |
| Y319L-F | AGGATTCCcttTACGGCGGCAACCCCTTGG |
| Y319L-R | GCCGTAaagGGAATCCTCGGGGTAGCGG |
| Y320H-F | AGGATTCCTACcacGGCGGCAACCCCTTGG |
| Y320H-R | GCCgtgGTAGGAATCCTCGGGGTAGCGG |
| S353T-F | ATCACCATCACCaccACCTCCCTCGCTTTCTTCAAGG |
| S353T-R | GTggtGGTGATGGTGATGGAGCCGATC |
| S365A-F | ATGTGTACCCCgctGCTGCGACGGGTACCTACG |
| S365A-R | AGCagcGGGGTACACATCCTTGAAGAAAGCGA |
| A367E-F | GCTgagACGGGTACCTACGCCTCTGG |
| A367E-R | TAGGTACCCGTctcAGCGGAGGGGTACACATCC |
| A372S-F | TACCTACtctTCTGGCAGCACTACCTTCAATG |
| A372S-R | TGCCAGAagaGTAGGTACCCGTCGCAGC |
| S373K-F | cgccactGGCAGCACTACCTTCAATGCCA |
| S373K-R | AAGGTAGTGCTGCCagtGGCGTAGGTACCCGTCGC |
| S383A-F | CATCATCgctGCTGTGAAGACTTATGCCGACG |
| S383A-R | TCACAGCagcGATGATGGCATTGAAGGTAGTGC |
| T413R-F | AATTTGACCGCcgcACGGGTCTCTCCATCTCCGC |
| T413R-R | CGTgcgGCGGTCAAATTGCTCGGAAAGAGA |
| T482G-F | TTTGACCAGCGGCggtGCTGCTCCTTCGTCTACGTCG |
| T482G-R | CaccGCCGCTGGTCAAAGTAGCGG |
| T513E-F | TTGCACCACCCCCgagGCCGTCGCTGTGACCTTCG |
| T513E-R | CctcGGGGGTGGTGCAAGAGCC |
| T526S-F | CACCACCtctTTTGGTGAGAATGTCTACCTGGTCGG |
| T526S-R | CACCAAAagaGGTGGTGGCAATCTCATCGAAG |
| F527T-F | ACCaccGGTGAGAATGTCTACCTGGTCGGTTC |
| F527T-R | ACATTCTCACCggtGGTGGTGGTGGCAATCTCAT |
| A553S-F | CTGAGCagcAGCAAGTACACTTCGAGCAACCC |
| A553S-R | TACTTGCTgctGCTCAGGGGAATACCGTTGGC |
| K555A-F | CGCTAGCgctTACACTTCGAGCAACCCTCTGTG |
| K555A-R | AAGTGTAagcGCTAGCGCTCAGGGGAATAC |
| A609S-F | CACTACTtccACCGAGAACGACACCTGGC |

| | |
|---|---|
| A609S-R | TCTCGGTggaAGTAGTGGTACCGCACTTGGC |

**Table S2.** Primers used for the construction of PoGA variants prioritized by the CASPEA model.

| Name | 5' --> 3' sequence |
|---|---|
| Q93E-F | AGTATGTGAACGCCgagGCTAAGCTCCAGACGGTTTCC |
| Q93E-R | ctcGGCGTTCACATACTGCTCAATGACG |
| S100N-F | GACGGTTaacAACCCTTCTGGAGGCCTCTCTG |
| S100N-R | AAGGGTTgttAACCGTCTGGAGCTTAGCCTGG |
| Q233G-F | CTACATGggcTCCTTCTGGACCGGCTCCTATA |
| Q233G-R | AGAAGGAgccCATGTAGCAGAGAATCTGAGGAGC |
| S234G-F | TACATGCAGggcTTCTGGACCGGCTCCTATATCA |
| S234G-R | CAGAAgccCTGCATGTAGCAGAGAATCTGAGG |
| N242D-F | CTATATCgacGCCAACACTGGCGGT |
| N242D-R | TGTTGGCgtcGATATAGGAGCCGGTCCAGAAGGA |
| A243G-F | TATCAACggcAACACTGGCGGTGGCCG |
| A243G-R | CAGTGTTgccGTTGATATAGGAGCCGGTCCAG |
| A304G-F | AACTCCGGAATTggcCAGGGCAAGGCCGTATCTG |
| A304G-R | TGgccAATTCCGGAGTTCAGAGCGTAGAC |
| Q305N-F | AATTGCCaacGGCAAGGCCGTATCTGTCGGCC |
| Q305N-R | CCTTGCCgttGGCAATTCCGGAGTTCAGAGCG |
| I304N-F | CGATGCCaacTATCAGTGGAACAAGATCGGCTC |
| I304N-R | ACTGATAgttGGCATCGTAGAGCTGCTCG |
| Y341F-F | TGCCATCttcCAGTGGAACAAGATCGGCTCCA |
| Y341F-R | TCCACTGgaaGATGGCATCGTAGAGCTGCTCG |
| A357K-F | CTCCCTCaagTTCTTCAAGGATGTGTACCCCTCCG |
| A357K-R | TGAAGAActtGAGGGAGGTGCTGGTGATGGTG |
| A380N-F | TaatATCATCAGCGCTGTGAAGACTTATGC |
| A380N-R | CAGCGCTGATGATattATTGAAGGTAGTGCTGCCAGAGG |
| I382E-F | TCAATGCCATCgagAGCGCTGTGAAGACTTATGCCG |
| I382E-R | GCTctcGATGGCATTGAAGGTAGTGCTGCCAG |
| S407P-F | CTCTCTTcccGAGCAATTTGACCGCACCACG |
| S407P-R | ATTGCTCgggAAGAGAGCCGTTGGCGTAGGAG |
| S417P-F | ACGGGTCTCcccATCTCCGCTCGTGACCTCA |
| S417P-R | GAGATgggGAGACCCGTGGTGCGGTCAA |
| I418N-F | TCTCTCCaacTCCGCTCGTGACCTCACCT |
| I418N-R | GAGCGGAgttGGAGAGACCCGTGGTGCG |
| S419A-F | TCTCCATCgccGCTCGTGACCTCACCTGG |
| S419F-R | ACGAGCggcGATGGAGAGACCCGTGGTGCG |
| A429G-F | CTACGCTggtCTCCTGACTGCCAACGACC |
| A429G-R | TCAGGAGaccAGCGTAGGACCAGGTGAGGTCA |
| R436N-F | TGACTGCCAACGACaacCGCAACGGTGTCGTCCC |
| R436N-R | gttGTCGTTGGCAGTCAGGAGTGCAG |
| N612V-F | TACCGAGgtcGACACCTGGCGCTAAGAATTCC |

| | |
|---|---|
| N612V-R | AGGTGTCgacCTCGGTAGCAGTAGTGGTACCGC |
| T614V-F | ACgtcTGGCGCTAAGAATTCCCTAGGG |
| T614V-R | TTCTTAGCGCCAgacGTCGTTCTCGGTAGCAGTAGTGG |

13. Rahban, M.; Zolghadri, S.; Salehi, N.; Ahmad, F.; Haertl, T.; Rezaei-Ghaleh, N.; Sawyer, L.; Saboury, A. A., Thermal stability enhancement: Fundamental concepts of protein engineering strategies to

manipulate the flexible structure. Int. J. Biol. Macromol. 2022, 214, 642–654.

14. Tan, K. P.; Varadarajan, R.; Madhusudhan, M. S., DEPTH: a web server to compute depth and predict small-molecule binding cavities in proteins. Nucleic Acids Res. 2011, 39, W242–W248.
